# Phylogenetic Association and Genetic Factors in Cold Stress Tolerance in *Campylobacter jejuni*

**DOI:** 10.1101/2022.08.13.503864

**Authors:** Jeong In Hur, Jinshil Kim, Sangryeol Ryu, Byeonghwa Jeon

## Abstract

*Campylobacter jejuni* is a major foodborne pathogen transmitted to humans primarily via contaminated poultry meat. Since poultry meat is generally processed, distributed, and stored in the cold chain, the survival of *C. jejuni* at refrigeration temperatures crucially affects human exposure to *C. jejuni*. Here, we investigated genetic factors associated with cold stress tolerance in *C. jejuni*. Seventy-nine *C. jejuni* strains isolated from retail raw chicken exhibited different survival levels at 4°C for 21 days. Multilocus sequence typing (MLST) clonal complex (CC)-21 and CC-443 were dominant among cold stress-tolerant strains, whereas CC-45 was common among cold stress-sensitive strains. Genome-wide average nucleotide identity (ANI) analysis identified a phylogenetic cluster associated with cold stress tolerance. Moreover, a pan-genome analysis revealed 58 genes distinctively present in the cold stress-tolerant phylogenetic cluster. Among the 58 genes, *cfrA*, encoding the ferric enterobactin receptor involved in ion transport and metabolism, was selected for further analysis. Remarkably, the viability of a Δ*cfrA* mutant at 4°C was significantly decreased, while the levels of total reactive oxygen species and intracellular iron exceeded those in the wild type. Additionally, a knockout mutation of *cfrA* also significantly decreased the viability of three cold stress-tolerant isolates at 4°C, confirming the role of *cfrA* in cold stress tolerance. The results of this study demonstrate that unique phylogenetic clusters of *C. jejuni* associated with cold stress tolerance exist and that *cfrA* is a genetic factor contributing to cold stress tolerance in *C. jejuni*.

**IMPORTANCE:** The tolerance of foodborne pathogens to environmental stresses significantly affects food safety. Several studies have demonstrated that *C. jejuni* survives extended exposure to low temperatures, but the mechanisms of cold stress tolerance are not fully understood. Here, we demonstrate that *C. jejuni* strains in certain phylogenetic groups exhibit increased tolerance to cold stress. Notably, *cfrA* is present in the phylogenetic cluster associated with cold stress tolerance and plays a role in *C. jejuni* survival at low temperatures by alleviating oxidative stress. This is the first study to discover phylogenetic associations involving cold stress tolerance and identify genetic elements conferring cold stress tolerance in *C. jejuni*.

## INTRODUCTION

*Campylobacter jejuni* is a major cause of acute gastroenteritis in humans (1-3). Human infection by *C. jejuni* is frequently associated with the consumption of contaminated poultry meat (4, 5), manifesting clinical symptoms such as diarrhea, abdominal cramps, and fever (6). In some cases, *C. jejuni* infection can result in Guillain–Barré syndrome, a neuropathy causing muscular paralysis, as a postinfection complication (7, 8). Food industries in most developed countries adopt cold-chain processing and distribution of meat products to ensure food safety and quality (3, 9). Although *C. jejuni*, as a thermotolerant species, can optimally grow at elevated temperatures, such as 42°C, the survival of *C. jejuni* on poultry meat in the cold chain poses a food safety threat (10, 11).

Most foodborne pathogens, such as *Bacillus, Salmonella*, and *Escherichia coli*, produce cold-shock proteins (12-14). When exposed to cold shock, *E. coli* increases the expression of cold-shock proteins, such as CspA (15, 16), which helps bacteria survive at low temperatures by disaggregating and reactivating proteins unfolded or misfolded by the temperature downshift (17, 18). As noted above, most human campylobacteriosis cases are primarily caused by the consumption of contaminated poultry. This suggests that despite the lack of cold-shock proteins, *C. jejuni* can successfully survive extended exposure to low temperatures of the cold chain during the distribution and storage of poultry products (11). Studies thus far show that oxidative stress defense is associated with cold stress tolerance in *Campylobacter* (18, 19). Exposure to cold stress increases the expression of oxidative stress defense genes in *C. jejuni* (19). Moreover, a knockout mutation of *sodB* encoding superoxide dismutase (SodB) compromises viability after freeze–thaw stress (20).

Iron is essential for various physiological processes; however, excessive iron disrupts redox homeostasis and catalyzes the generation of reactive oxygen species (ROS) via the Fenton reaction under stress conditions (21-23). ROS cause oxidative damage to cellular components, such as DNA and proteins, and can lead to cell death (24). Since the iron-catalyzed Fenton reaction converts hydrogen peroxide to hydroxyl radicals, the most noxious ROS causing cellular damage, intracellular free iron levels can be correlated with oxidative stress (23). The expression of iron-related genes is elevated in *C. jejuni* during cold shock (19), suggesting that iron may play an essential role in the adaptation of *C. jejuni* to cold shock. However, little is understood about how *C. jejuni* can tolerate low temperatures of the cold chain during foodborne transmission to humans via refrigerated poultry meat.

To fill this knowledge gap, in this study, we investigated cold tolerance in 79 *C. jejuni* strains isolated from retail raw chicken in our previous study (25) and discovered that some strains of *C. jejuni* are highly tolerant to cold stress. Moreover, cold stress tolerance is associated with specific clonal complexes (CCs), which indicates that strains with cold stress tolerance are phylogenetically related. By comparing 79 *C. jejuni* isolates and testing them with gene knockout mutations, we show that *cfrA* contributes to cold stress tolerance in *C. jejuni*. Notably, we demonstrate that intracellular iron and oxidative stress defense are related to cold stress tolerance driven by *cfrA* in *C. jejuni*.

## RESULTS

### Phylogenetic association with cold stress tolerance in *C. jejuni*

*C. jejuni* can be isolated from refrigerated poultry meat and various environmental samples from poultry farms, although it is a thermotolerant species (26, 27). Thus, we hypothesized that *C. jejuni* circulating in poultry production may have the capability to tolerate cold temperatures. Using 79 *C. jejuni* strains isolated from retail raw chicken in our previous study (25), we first measured the survival of *C. jejuni* at refrigeration temperature for 21 days. As the tested strains showed a wide range of viability at 4°C, we divided the 79 strains into two groups of equal size by their viability at 21 days and designated them as cold stress-tolerant (n=39) and cold stress-sensitive (n=40), respectively. The viable counts of the cold stress-tolerant strains at 4°C at all sampling times (7, 14, and 21 days) were significantly different from those of the cold stress-sensitive strains (Fig. 1A).

**Figure 1.**
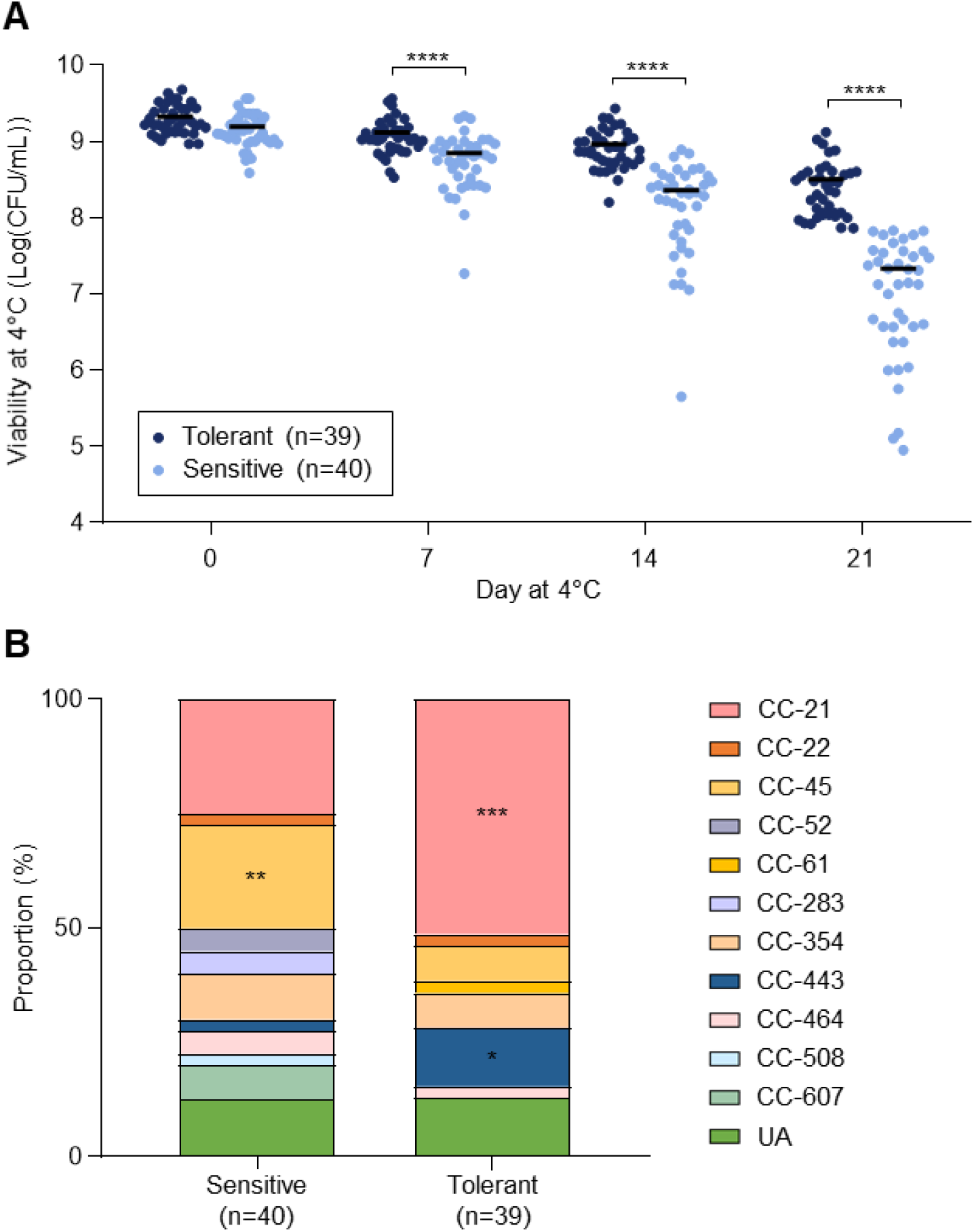
Different levels of cold stress tolerance in *C. jejuni* isolates from retail chicken and differences in proportions of MLST CCs between cold stress-sensitive and cold stress-tolerant strains. (A) Viable counts of 79 *C. jejuni* strains were measured at 0, 7, 14, and 21 days of exposure at 4°C. The Student’s *t* test was performed to compare viability between cold stress-sensitive and cold stress-tolerant strains. A solid black bar indicates the mean. The data are representative of three independent experiments showing similar results. (B) MLST CCs of the cold stress-sensitive and cold stress-tolerant strains were grouped based on viability after cold stress exposure for 21 days. A chi-square test was conducted to determine whether the CC proportions were statistically related to cold stress tolerance. ^*^, *P* < 0.05; ^**^, *P* < 0.01; ^***^, *P* < 0.001; ^****^, *P* < 0.0001; CC, clonal complex; UA, unassigned to any CC defined.

To determine whether cold stress tolerance is related to bacterial phylogeny in *C. jejuni*, we compared multilocus sequence typing (MLST) CCs between cold stress-tolerant and cold stress-sensitive strains. Notably, CC-21 and CC-443 were predominant in cold stress-tolerant strains (51.3% and 12.8%, respectively), whereas CC-45 was dominant (22.5%) in cold stress-sensitive strains (Fig. 1B). The associations of CC-21, CC-45, and CC-443 with cold stress tolerance were statistically significant (Fig. S1). These findings demonstrate that some *C. jejuni* strains are highly tolerant to low temperatures and that cold stress tolerance is phylogenetically associated in *C. jejuni*. These data also suggest that genetic elements involved in cold stress tolerance may exist in *C. jejuni*.

### *C. jejuni* strains tolerant or sensitive to cold stress are phylogenetically distinct

We performed genome-wide average nucleotide identity (ANI) analysis to further investigate the phylogenetic association with cold stress tolerance. As a result, we identified four phylogenetic clusters that are distinctly separate below the 98% ANI threshold: Cluster 1 (n=6), Cluster 2 (n=15), Cluster 3 (n=26), and Cluster 4 (n=32) (Fig. 2). Interestingly, the phylogenetic clusters were related to MLST CCs and cold stress tolerance. When MLST CCs were compared, Cluster 2 showed a significantly high proportion of CC-45, while CC-443 and CC-21 were highly prevalent in Cluster 3 and Cluster 4, respectively (Fig. 3A). Consistent with the patterns of cold stress tolerance of these CCs (Fig. 1B), Cluster 2 and Cluster 4 consisted mostly of cold stress-sensitive and cold stress-tolerant strains, respectively (Fig. S2). Moreover, the viability of *C. jejuni* after 21 days of exposure to cold stress was significantly different between Cluster 2 and Cluster 4 (Fig. 3B). These results suggest that cold stress tolerance is associated with genetic backgrounds in *C. jejuni*.

**Figure 2.**
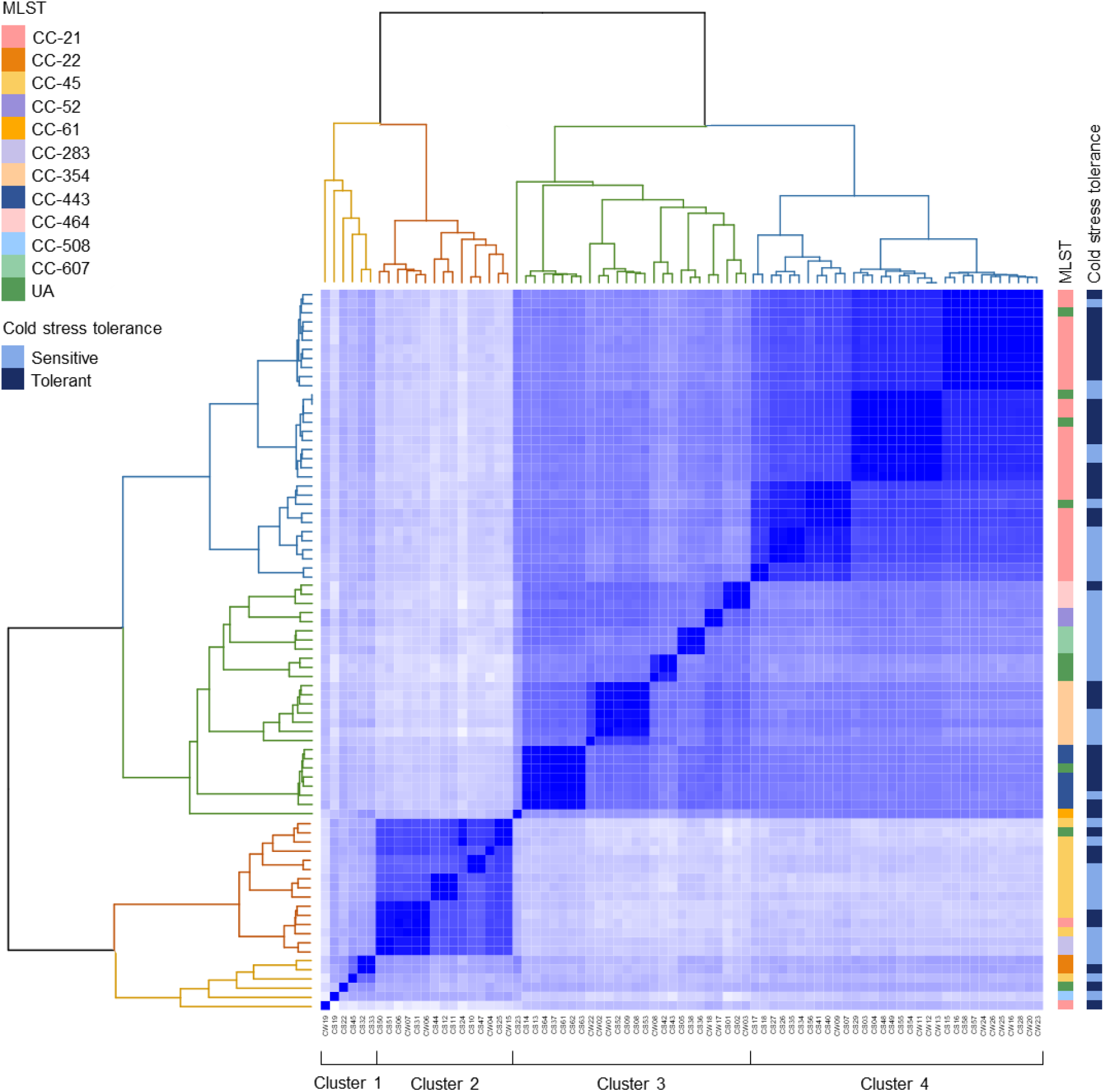
Identification of phylogenetic clusters related to cold stress tolerance in *C. jejuni*. Genome-wide ANI values separated 79 *C. jejuni* strains into four clusters: Cluster 1 (yellow, n=6), Cluster 2 (orange, n=15), Cluster 3 (green, n=26), and Cluster 4 (blue, n=32). The information about MLST and cold stress tolerance of 79 *C. jejuni* strains is indicated on the right side of the heatmap. CC, clonal complex; UA, unassigned to any CC defined.

**Figure 3.**
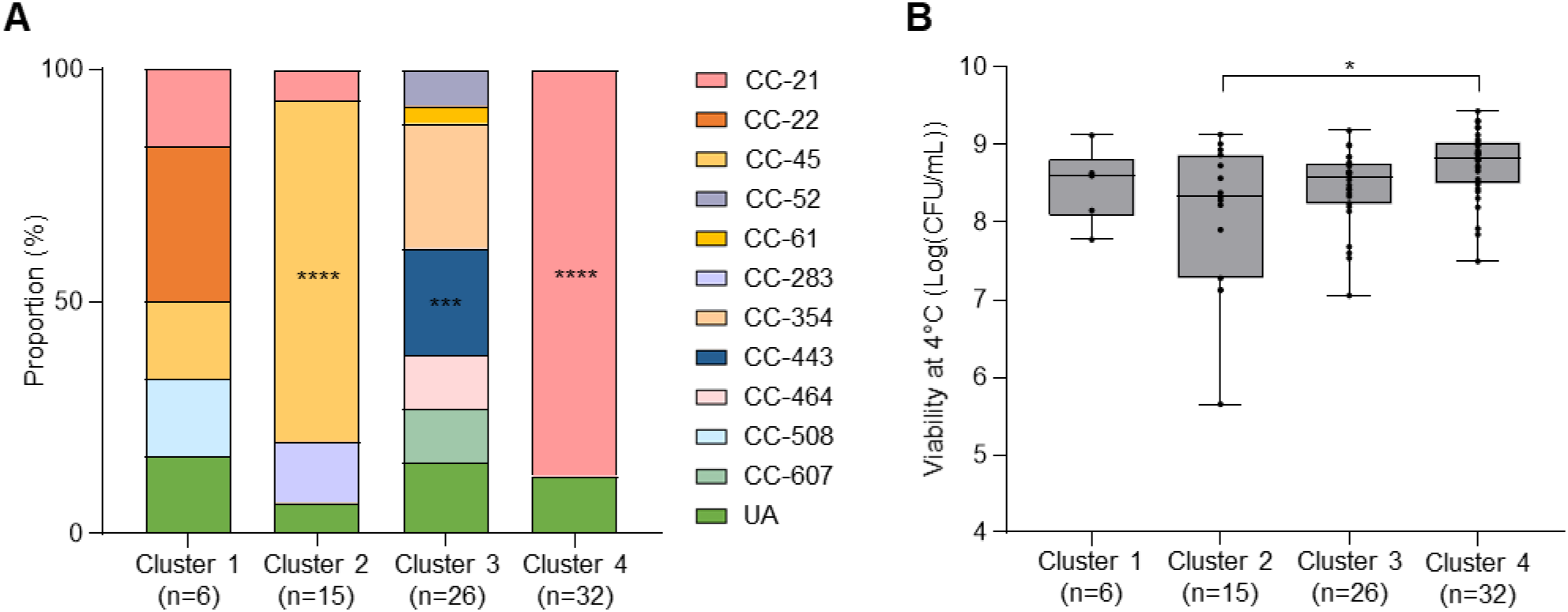
Association of phylogenetic clusters with MLST CCs and cold stress tolerance. (A) Proportions of MLST CCs in the four phylogenetic clusters. A chi-square test was conducted for statistical analysis. (B) Viability at 4°C for 21 days of *C. jejuni* strains of the four phylogenetic clusters. The results indicate means and standard deviations. The Student’s *t* test was performed to compare viability between the two clusters. ^*^, *P* < 0.05; ^***^, *P* < 0.001; ^****^, *P* < 0.0001; CC, clonal complex; UA, unassigned to any CC defined.

### Genetic elements are unique to cold stress-tolerant strains of *C. jejuni*

Cluster 2 and Cluster 4 were phylogenetically distant (Fig. 2) and showed significantly different levels of cold stress tolerance (Fig. 3B). Thus, we conducted a pan-genome analysis to identify genes potentially associated with cold stress tolerance by comparing the two clusters. The analysis revealed 58 genes that are present in the cold-tolerant cluster (i.e., Cluster 4) and absent from the cold-sensitive cluster (i.e., Cluster 2) (Fig. S3, Table 1). The 58 genes are involved in various functions, including inorganic ion transport and metabolism, amino acid transport and metabolism, defense mechanisms, transcription, and carbohydrate transport and metabolism (Table 1). Among the 58 genes discovered by comparing Cluster 2 and Cluster 4, we decided to investigate how iron metabolism genes can be involved in cold stress tolerance and selected *cfrA* encoding the ferric enterobactin receptor for further investigation in the remainder of the study. When we examined the occurrence of *cfrA* in the 79 *C. jejuni* isolates, there was a clear separation of phylogenetic groups (Fig. 4A). The strains lacking *cfrA* belonged predominantly to CC-45 (60.0%), the CC associated with cold stress sensitivity, and the strains harboring *cfrA* belonged predominantly to CC-21 (47.5%), UA (15.3%), CC-354 (11.9%), and CC-443 (10.2%) (Fig. 4B). Notably, these results are consistent with the results of viability assays and ANI analysis, which also show that strains in CC-45 are related to cold stress sensitivity and that those in CC-21 and CC-443 tend to be tolerant to cold stress (Figs. 1 and 2). These data suggest that *cfrA* can be involved in cold stress tolerance in *C. jejuni*.

**Table 1.** Fifty-eight genes present in the cold-tolerant cluster and absent in the cold-sensitive cluster.

| Locus tag | Gene | Description |
| --- | --- | --- |
| Inorganic ion transport and metabolism |  |  |
| *cj0444* | pseudogene | TonB-dependent receptor |
| *cj0676* | *kdpA* | potassium-transporting ATPase subunit KdpA |
| *cj0755* | *cfrA* | ferric enterobactin receptor CfrA |
| *cj1040c* |  | MFS transporter |
| *cj1415c* | *cysC* | adenylyl-sulfate kinase |
| Amino acid transport and metabolism |  |  |
| *cj0029* | *ansA* | type II asparaginase |
| *cj0481* | *dapA* | dihydrodipicolinate synthase family protein |
| *cj0763c* | *hisS* | histidine--tRNA ligase |
| *cj0817* | *glnH* | transporter substrate-binding domain-containing protein |
| *cj1726c* | *metA* | homoserine O-succinyltransferase |
| *cj1727c* | *metB* | O-acetylhomoserine aminocarboxypropyltransferase/cysteine synthase |
| Defense mechanisms |  |  |
| *cj0139* |  | hypothetical protein |
| *cj0690c* |  | SAM-dependent DNA methyltransferase |
| *cj1549c* | *hsdR* | type I restriction endonuclease subunit R |
| *cj1551c* | *hsdS* | restriction endonuclease subunit S |
| *cj1553c* | *hsdM* | SAM-dependent DNA methyltransferase |
| Transcription |  |  |
| *cj0480c* |  | IcIR family transcriptional regulator |
| *cj0757* | *hrcA* | HrcA family transcriptional regulator |
| *cj1552c* | *mloB* | transcriptional regulator |
| *cj1556* |  | helix-turn-helix transcriptional regulator |
| Carbohydrate transport and metabolism |  |  |
| *cj0482* | *uxaA'* | UxaA family hydrolase |
| *cj0483* | *uxaA'* | UxaA family hydrolase |
| *cj0484* |  | MFS transporter |
| Secondary metabolites biosynthesis, transport and catabolism |  |  |
| *cj0170* |  | methyltransferase domain-containing protein |
| *cj1325* |  | methyltransferase domain-containing protein |
| *cj1420c* |  | class I SAM-dependent methyltransferase |
| Energy production and conversion |  |  |
| *cj0490* | *ald'* | aldehyde dehydrogenase |
| *cj1585c* |  | FAD-binding oxidoreductase |
| Nucleotide transport and metabolism |  |  |
| *cj0766* | pseudogene | Putative arylsulfate sulfotransferase |
| *cj0381c* | *pyrF* | orotidine-5'-phosphate decarboxylase |
| General function prediction only |  |  |
| *cj0054c* |  | TIGR00730 family Rossman fold protein |
| *cj1555c* |  | NAD(P)-dependent oxidoreductase |
| Lipid transport and metabolism |  |  |
| *cj0485* |  | SDR family oxidoreductase |
| Posttranslational modification, protein turnover, chaperones |  |  |
| *cj1725* |  | NAD(P)/FAD-dependent oxidoreductase |
| Intracellular trafficking, secretion, and vesicular transport |  |  |
| *cj0969* | pseudogene | hemagglutination domain protein |
| Function unknown |  |  |
| *cj0055c* |  | hypothetical protein |
| *cj0056c* |  | hypothetical protein |
| *cj0247c* |  | chemotaxis protein |
| *cj0380c* |  | hypothetical protein |
| *cj0425* |  | hypothetical protein |
| *cj0565* |  | hypothetical protein |
| *cj0566* |  | hypothetical protein |
| *cj0568* |  | hypothetical protein |
| *cj0569* |  | hypothetical protein |
| *cj0617* |  | DUF2920 family protein |
| *cj0685c* | *cipA* | DUF2972 domain-containing protein |
| *cj0740* |  | hypothetical protein |
| *cj0818* |  | hypothetical protein |
| *cj0859c* |  | hypothetical protein |
| *cj0866* | pseudogene | arylsulfate sulfotransferase |
| *cj0970* |  | hypothetical protein |
| *cj0972* |  | hypothetical protein |
| *cj0978c* |  | putative lipoprotein |
| *cj1122c* |  | hypothetical protein |
| *cj1456c* |  | hypothetical protein |
| *cj1550c* | *rloH* | AAA family ATPase |
| *cj1558* |  | Permease |
| *cj1714* |  | small hydrophobic protein |

**Figure 4.**
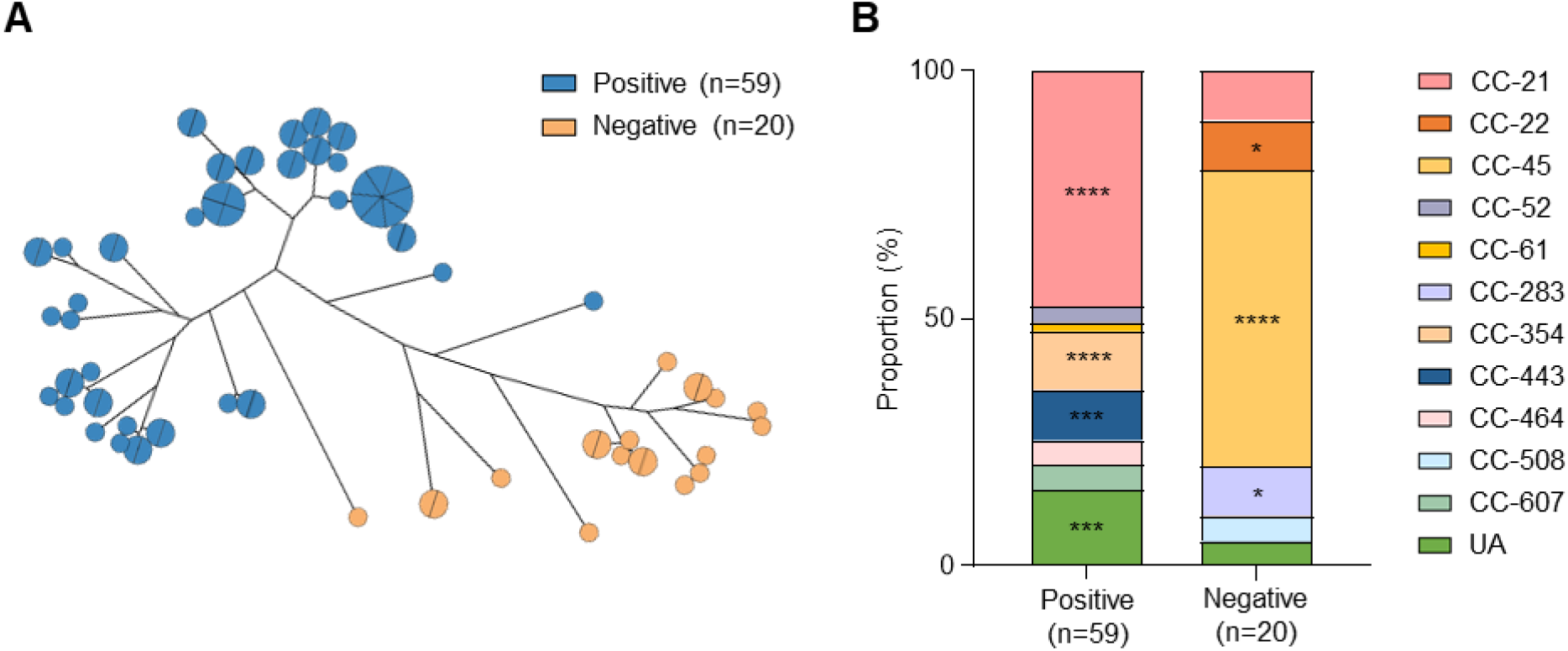
Phylogenetic distinction and MLST CC composition depending on the presence of *cfrA*. (A) The minimum spanning tree was generated by core gene alignment obtained from pan-genome analysis. (B) MLST CCs of 79 *C. jejuni* strains were compared between *cfrA*-positive and *cfrA*-negative phylogenies. A chi-square test was conducted for comparison of the proportions of the CCs. ^*^, *P* < 0.05; ^***^, *P* < 0.001; ^****^, *P* < 0.0001; CC, clonal complex; UA, unassigned to any CC defined.

### CfrA contributes to the survival of *C. jejuni* at refrigeration temperatures

Studies show that a *sodB* mutation compromises the survival of *C. jejuni* at refrigeration temperatures, indicating that oxidative stress defense is related to cold stress tolerance (18, 19). Intracellular iron affects oxidative stress through the Fenton chemistry (28, 29). Therefore, we hypothesized that *cfrA* may be involved in cold stress tolerance by affecting oxidative stress in *C. jejuni* at refrigeration temperatures. Before testing the hypothesis using the Δ*cfrA* mutant, we questioned whether cold stress could induce oxidative stress in *C. jejuni*. We observed that exposure to cold stress at 4°C led to total ROS accumulation in *C. jejuni* (Fig. 5). The levels of total ROS accumulation at 4°C were similar under microaerobic and aerobic conditions (Fig. 5). These results suggest that *C. jejuni* under cold stress conditions experiences increased oxidative stress at a level similar to that under aerobic conditions.

**Figure 5.**
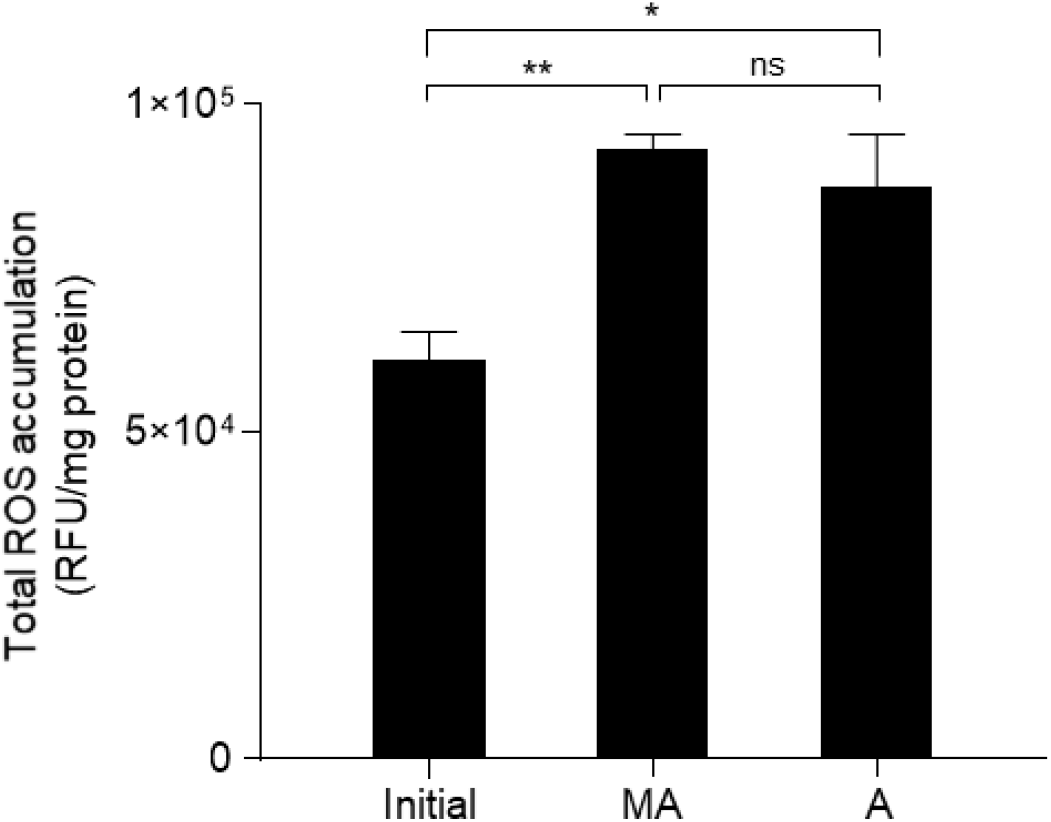
Increased oxidative stress after exposure to cold stress in *C. jejuni* under microaerobic and aerobic conditions. Total ROS accumulation levels were measured before (Initial) and after exposure to cold stress for 4 days under microaerobic (MA) or aerobic (A) conditions. The experiment was repeated three times. Each bar indicates the standard error of the means. The Student’s *t* test was performed for statistical analysis. ^*^, *P* < 0.05; ^**^, *P* < 0.01; ns, non-significant; RFU, relative fluorescence units.

To further examine whether *cfrA* is involved in cold stress tolerance, we constructed a Δ*cfrA* mutant. In addition to genetic confirmation of a mutation by sequencing (data not shown), the mutation was confirmed phenotypically by observing a defect in the uptake of the ferric enterobactin complex in the Δ*cfrA* mutant (Fig. 6A). Remarkably, the viability of a Δ*cfrA* mutant at 4°C was significantly decreased compared to that of the wild type (WT) (Fig. 6B). Genetic complementation of the Δ*cfrA* mutant with an intact copy of *cfrA* fully restored cold stress tolerance to the WT level (Fig. 6B). Since *cfrA* is related to iron metabolism, we measured the intracellular iron level before and after exposure to cold stress. Interestingly, a Δ*cfrA* mutation significantly elevated the iron level (Fig. 6C). These results suggest that *cfrA* is associated with the control of intracellular iron levels in *C. jejuni* under cold stress conditions. Moreover, exposure to cold stress significantly increased ROS levels in the Δ*cfrA* mutant compared to WT (Fig. 6D). These results suggest that *C. jejuni* confronts increased oxidative stress at cold temperatures and that *cfrA* contributes to cold stress tolerance by controlling intracellular iron and oxidative stress. Finally, we confirmed the role of *cfrA* in cold stress tolerance using three cold stress-tolerant isolates. The three strains were selected from CCs that comprise large proportions among cold stress-tolerant strains: CS14 (CC-443), CS49 (CC-21), and CS62 (CC-443). We constructed Δ*cfrA* mutants of the three cold stress-tolerant strains to validate the role of *cfrA* in cold stress-tolerant strains. Notably, a knockout mutation of *cfrA* significantly compromised the viability of the three cold stress-tolerant strains of *C. jejuni* at 4°C when compared to their WTs (Fig. 6E). These data suggest that *cfrA* contributes to cold stress tolerance in *C. jejuni* by alleviating oxidative stress.

**Figure 6.**
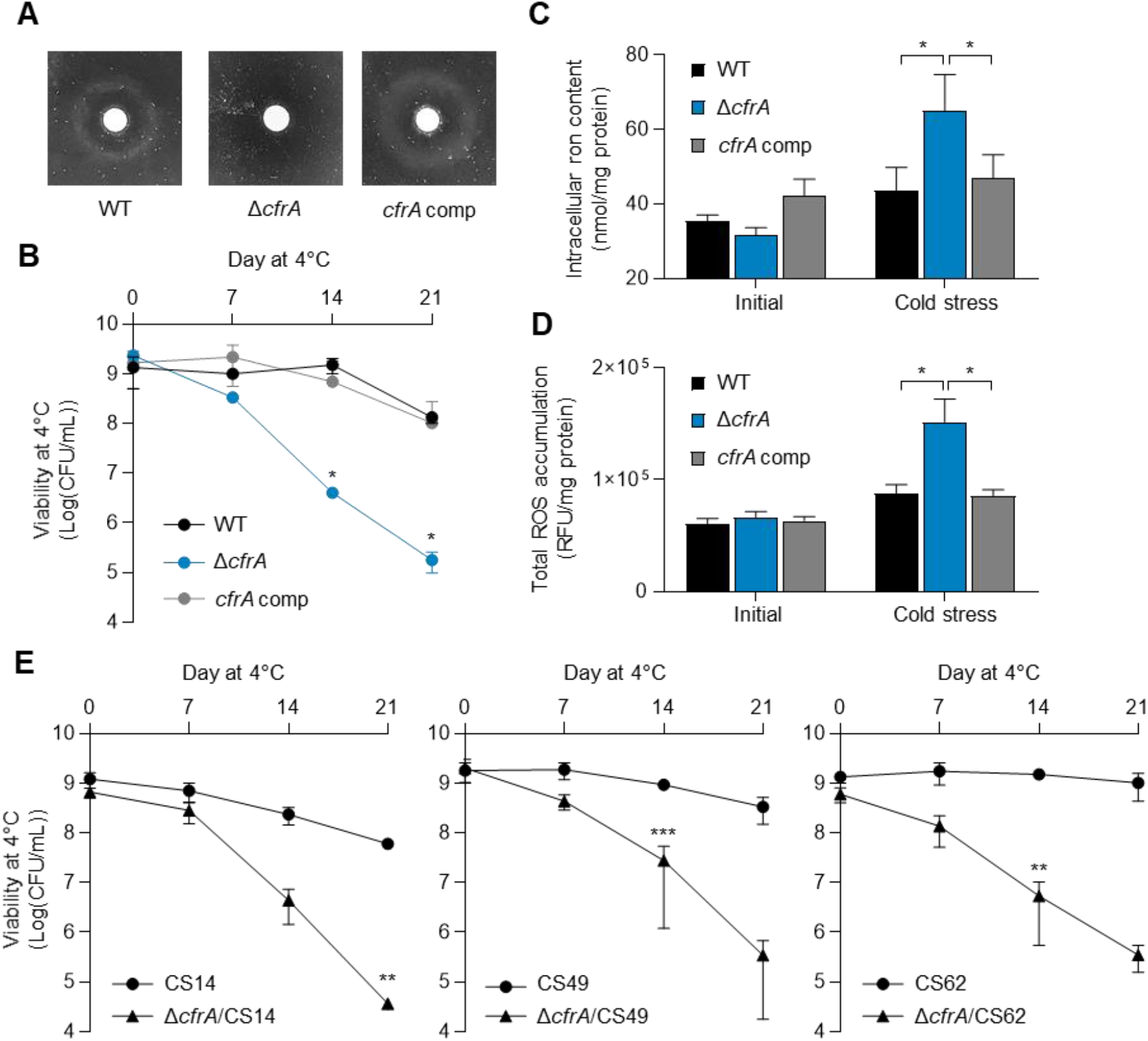
Contribution of *cfrA* to cold stress tolerance in *C. jejuni*. (A) The inability of a Δ*cfrA* mutant to take up enterobactin. (B) Defective cold stress tolerance in the Δ*cfrA* mutant. The asterisk indicates a significant difference in viability between the Δ*cfrA* mutant and WT at the same sampling time. (C) Intracellular iron levels in *C. jejuni* before and after exposure to cold stress at 4°C for 4 days. (D) Total ROS accumulation in *C. jejuni* before and after exposure to cold stress for 4 days. (E) Significant defects in cold stress tolerance in three cold stress-tolerant strains of *C. jejuni*. The asterisks indicate the statistical significance of differences in viability between the Δ*cfrA* mutant and wild type at the same sampling time after exposure to cold stress. The experiment was repeated three times and produced similar results. The error bars show the standard errors of the means. The Student’s *t* test was performed for statistical analysis. ^*^, *P* < 0.05; ^**^, *P* < 0.01; ^***^, *P* < 0.001; WT, *C. jejuni* NCTC 11168 wild type; Δ*cfrA*, Δ*cfrA* mutant; *cfrA* comp, *cfrA*-complemented strain.

## DISCUSSION

*C. jejuni* is a major foodborne pathogen transmitted to humans via contaminated poultry meat. Considering the use of the cold chain to process and distribute poultry meat, cold stress is one of the major stress conditions *C. jejuni* must overcome during foodborne transmission to humans. However, little attention has been given to cold stress tolerance in *C. jejuni*. Here, we tested cold stress tolerance in 79 *C. jejuni* strains isolated from retail raw chicken and discovered that the level of cold stress tolerance varies in *C. jejuni* depending on the strain (Fig. 1A). Moreover, strains in CC-21 and CC-443 were significantly more likely to show cold stress tolerance, and those in CC-45 were more likely to exhibit cold stress sensitivity (Fig. 1B). A previous study also showed that *C. jejuni* strains belonging to CC-21 survived better at 4°C than those in CC-45 (31). Phylogenetic studies demonstrate that CC-21 and CC-443 are closely related to each other, whereas CC-45 is more distant (32, 33). CC-21 and CC-45 are the major generalist CCs occupying the diverse population of *C. jejuni* isolated from multiple different hosts, such as chickens, cattle, and wild birds (34-36). CC-443 is frequently associated with chickens (33). An MLST analysis of 1,215 isolates from human campylobacteriosis cases in New Zealand over nine years showed that CC-45 is characteristic in summer, while CC-21 peaks in late autumn to early winter, exhibiting the seasonal prevalence of *C. jejuni* strains belonging to CC-21 and CC-45 (37). A similar pattern of summer seasonality of CC-45 has also been reported in the UK (30). Based on the association of CC-45 with cold stress sensitivity revealed in this study (Fig. 1B), it can be speculated that *C. jejuni* strains belonging to CC-45 may be less prevalent in poultry production environments in winter and may cause human infections primarily in summer.

The phylogenetic analysis using whole-genome sequences divided the 79 strains into four clusters based on ANI analysis (Fig. 2) and identified two clusters associated with cold stress tolerance (Fig. 3). Comparing the genome sequences between the two clusters led to the identification of 58 genes present in *C. jejuni* strains in the cold stress-tolerant cluster and absent from the strains in the cold stress-sensitive cluster (Table 1). Based on previous efforts to identify genes involved in human campylobacteriosis (38, 39), interestingly, most of the 58 genes unique to the cold stress-tolerant cluster are present only in clinical isolates and absent from nonclinical isolates, including *kpsA* (encoding potassium-transporting ATPase subunit), *uxaA* (encoding UxaA family hydrolase), *cfrA* (encoding ferric enterobactin receptor) and others. Although it remains unexplained whether these genes are related to the pathogenicity of *C. jejuni*, it can be speculated that cold stress-tolerant strains are more likely to cause human infection than cold stress-sensitive strains because cold tolerance enables *C. jejuni* to survive on poultry meat, the primary cause of campylobacteriosis, in the food supply chain and increases the chances of human exposure to *C. jejuni*.

Notably, our findings demonstrate that *cfrA* plays a role in cold stress tolerance in *C. jejuni*. In a previous study, genes related to iron metabolism, including *cfrA*, were found to be crucial for bacterial survival under stressful conditions during host colonization (40). The phylogenetic analysis of core gene alignment shows a clear distinction between *cfrA*-positive and *cfrA*-negative phylogenies (Fig. 4A). CC-21 & CC-443 and CC-45, which are correlated with cold stress tolerance and sensitivity, respectively, are separated based on the presence of *cfrA* (Fig. 4B). Remarkably, the viability of the Δ*cfrA* mutant at 4°C was significantly compromised compared to that of WT (Fig. 6B). A Δ*cfrA* mutation also reduced cold stress tolerance in cold stress-tolerant isolates (Fig. 6E). Altogether, these results are the first to present the role of CfrA in cold stress tolerance in *C. jejuni*.

Studies have shown the association of oxidative stress defense with cold stress tolerance in *C. jejuni*. In particular, a mutation of *sodB* encoding superoxide dismutase makes *Campylobacter* susceptible to freeze–thaw stress (20, 41). We found that oxidative stress increases when *C. jejuni* is exposed to refrigeration temperatures (Fig. 6D). The levels of total ROS accumulation were similar between microaerobic and aerobic conditions (Fig. 5), indicating oxidative stress increases in *C. jejuni* at refrigeration temperatures regardless of oxygen level. A similar observation has been reported in another bacterium, where growth at 4°C increased oxidative stress and generated ROS in *Pseudomonas fluorescens* MTCC 667, an isolate from Antarctica (42). Presumably, reduced carbon metabolism at low temperatures may decrease the formation of the reducing compounds NADH and FADH_2_, subsequently increasing oxidative stress. In addition, we also observed that exposure to refrigeration temperatures increased the intracellular level of iron in *C. jejuni* (Fig. 6C). A knockout mutation of *cfrA* increased the levels of iron and total ROS (Fig. 6C and D), which may decrease viability at refrigeration temperatures because an iron upshift leads to oxidative stress and can trigger cell death. These data suggest that CfrA contributes to the control of intracellular iron and redox homeostasis in *C. jejuni* at refrigeration temperatures.

In summary, we demonstrated for the first time the phylogenetic association with cold stress tolerance in *C. jejuni* and showed that specific CCs are associated with cold stress tolerance. We also identified genes unique to the cold stress-tolerant cluster. Finally, we revealed that *cfrA* contributes to cold stress tolerance by controlling intracellular iron levels and oxidative stress. Future studies will need to elucidate the molecular mechanisms of cold stress tolerance driven by *cfrA*.

## MATERIALS AND METHODS

### Bacterial strains and culture conditions

Seventy-nine *Campylobacter jejuni* strains previously isolated from retail raw chicken were used in this study (25). *C. jejuni* NCTC 11168 was used as a reference strain in this study. The *C. jejuni* strains were routinely grown on Mueller–Hinton (MH) agar (Oxoid, Hampshire, UK) at 42°C for 18-24 h under microaerobic conditions (85% N_2_, 5% O_2,_ and 10% CO_2_) generated by Anoxomat (Mart Microbiology BV, Lichtenvoorde, The Netherlands).

### Cold stress tolerance test of *C. jejuni*

The survival of *C. jejuni* at 4°C was measured as described previously (43), with slight modifications. Briefly, an overnight culture on MH agar was resuspended in MH broth to an optical density of 600 nm (OD_600_) of 0.1 (ca, 10^9^ CFU/mL). The bacterial suspension was transferred to multiple 96-well plates in 200μL aliquots. To prevent sample desiccation, outer wells were filled with an equal volume of distilled water, and a container with water was placed near the 96-well plates. Wooden sticks were placed under both sides of the lids of the 96-well plates to improve air circulation. The 96-well plates were incubated at 4°C under aerobic conditions, and samples were taken after 0, 7, 14, and 21 days for serial dilution and bacterial counting. The strains with viable cells greater than 7.0 × 10^7^ CFU/mL after 21 days were called cold stress-tolerant strains, while those with viable cells less than 7.0 × 10^7^ CFU/mL after 21 days were called cold stress-sensitive strains.

### Whole-genome sequencing

Genomic DNA (gDNA) was extracted using a NucleoSpin Microbial DNA kit (Macherey-Nagel, PA, USA) and TissueLyser II (Qiagen, Hilden, Germany) according to the manufacturer’s instructions. A NanoDrop spectrophotometer (Thermo Fisher Scientific, OH, USA), gel electrophoresis, and Qubit Fluorometer (Thermo Fisher Scientific, OH, USA) were used to evaluate the quality of the gDNA. After the quality control of gDNA, the DNA library was prepared using the TruSeq Nano DNA LT Library Prep Kit (Illumina, CA, USA) according to the TruSeq Nano DNA Library Preparation protocol. The quality of the libraries was assessed on a 2100 Bioanalyzer System with a DNA1000 Chip (Agilent Technologies, CA, USA). Then, the constructed DNA libraries were sequenced with a 2 × 150 bp read length using the NextSeq 500 Sequencing System (Illumina, CA, USA).

### Bioinformatics analysis

Trimming and *de novo* assembly of raw reads generated from the whole genome sequencing were performed using CLC Genomic Workbench v20 with default parameters. Then, the assembled genomes were annotated using Prokka v1.14.6 with default parameters. To specify the degree of overall relatedness among genomes, we estimated the genome-wide ANI using FastANI v1.33. ANI estimates the average nucleotide identity of all orthologous genes shared between any two genomes. Organisms belonging to the same species typically exhibit 95% or higher ANI. Pairwise ANI values were visualized using a heatmap generated by ComplexHeatmap v2.2.0 and gplots v3.3.5 in R, dividing the strains into four phylogenetic clusters. In a search for characteristic genes present in the cold-tolerant cluster, pan-genome analysis was performed with Roary v3.11.2. For comparative analyses of the presence or absence of *cfrA*, minimum spanning trees were generated and visualized in GrapeTree v1.5.0 with core genome alignment obtained from Roary. Only the strains for which the presence or absence of *cfrA* was confirmed by PCR were used for the analysis described above. The primer sets are listed in Table S1.

### Construction of Δ*cfrA* mutants and a *cfrA*-complemented strain

A suicide plasmid carrying *cfrA* was constructed as described previously (44). Briefly, *cfrA* and its flanking region were amplified by PCR with GXL polymerase (Takara, Tokyo, Japan) from *C. jejuni* with the primers presented in Table S1. After digestion with *Sal*I and *Bam*HI, the PCR products were each ligated to pUC19 that had been treated with the same enzymes. The pUC19 plasmid containing *cfrA* was amplified by PCR from inside the gene with inverse primers using the same polymerase and ligated to a kanamycin cassette from pMW10. The suicide vectors were commercially sequenced by Bionics (Seoul, South Korea). These three plasmids were used as suicide vectors, and each vector was introduced into WT by electroporation. The *C. jejuni* culture was grown on MH agar plates containing kanamycin (50 μg/mL) to screen for Δ*cfrA* mutants. The *cfrA* mutation was confirmed by PCR and sequencing.

The complementation strain was constructed as previously described (45). Briefly, DNA fragments containing the intact copy of *cfrA* were amplified with primer pairs and cloned into a *Not*I site on a pUC19 derivative carrying an rRNA gene cluster (46, 47). Plasmids carrying *cfrA* were sequenced by Bionics (Seoul, South Korea) and used as complementation vectors. The complementation vectors were introduced into *cfrA* knockout mutants by electroporation. To screen for *cfrA* complementation strains, the *Campylobacter* culture was grown on MH agar plates containing kanamycin (50 μg/mL) and chloramphenicol (12.5 μg/mL). Complementation of *cfrA* was confirmed by PCR and sequencing.

### Measurement of ROS levels

ROS levels were measured as described previously with slight modifications (48). Total ROS accumulation level was measured using the fluorescent dye CM-H_2_DCFDA (Thermo Fisher Scientific, OH, USA). *C. jejuni* was prepared by an overnight culture on MH agar and resuspended in MH broth to an OD_600_ of 0.1. The bacterial suspension was transferred to a disposable culture tube (Kimble, NJ, USA) and incubated at 4°C. Samples were taken before and after exposure to cold stress for 4 days. After treatment with 10 μM CM-H_2_DCFDA for 30 min at room temperature, fluorescence was measured with a SpectraMax i3 Platform (Molecular Devices, CA, USA) at 495 nm excitation and 527 nm emission wavelengths. The fluorescence levels were normalized to the protein amounts determined with the Bradford assay (Bio-Rad, CA, USA).

### Growth promotion assay

As previous studies demonstrated that *Campylobacter* used ferric enterobactin as a sole source of iron during growth promotion assays (49), we measured the growth of *C. jejuni* strains as previously described (50). Briefly, an overnight culture on MH agar was resuspended in MH broth to an OD_600_ of 0.1. *C. jejuni* cells were grown in a disposable glass tube to log-phase. Deferoxamine mesylate salt (DFO) (Sigma Aldrich, MO, USA), a chelator, was added to melted MH agar at a final concentration of 20 μM. The cells were mixed with DFO-containing MH agar and adjusted to approximately 10^7^ CFU/mL. Each sample mixture was poured into Petri dishes for solidification. A sterile disk containing 25 μL of enterobactin (2 mM) (Sigma Aldrich, MO, USA) was placed on the surface of the agar in each dish. Autoclaved distilled water was used instead of enterobactin as a negative control.

### Measurement of intracellular iron levels

Levels of intracellular iron were measured as described previously (51). An overnight culture on MH agar was resuspended in MH broth to an OD_600_ of 0.1. *C. jejuni* cells were transferred to disposable culture tubes (Kimble, NJ, USA) in 3-mL aliquots and incubated at 4°C. Samples were taken before and after exposure to cold stress for 4 days. Briefly, the samples were washed twice with ice-cold PBS and disrupted with a sonicator. A standard curve was obtained by diluting 1 mM FeCl_3_ (Sigma Aldrich, MO, USA) standard solution. The samples were mixed with an iron-detection reagent (6.5 mM ferrozine, 6.5 mM neocuproine, 2.5 M ammonium acetate, and 1 M ascorbic acid) and incubated at room temperature for 30 min. The absorbance was measured with a SpectraMax i3 Platform (Molecular Devices, CA, USA) at 550 nm. The intracellular iron levels were normalized to the protein concentrations determined with Bradford assay (Bio-Rad, CA, USA).

### Statistical analysis

A chi-square test was performed when comparing proportions. The Student’s *t* test was performed for comparative analysis between two groups. GraphPad Prism (Version 8.0.1; GraphPad Software, Inc., CA, USA) was used for statistical analysis.

### Data availability

The GenBank accession numbers for the genome sequences of all 79 *C. jejuni* isolates used in the study are presented in Table S2.

## ACKNOWLEDGMENTS

J.I.H. and J.K. were supported by the BK21 Plus Program of the Department of Agricultural Biotechnology, Seoul National University, Seoul, Korea. This research was supported by a grant 19162MFDS037 from Ministry of Food and Drug Safety in 2021. This research was supported by Basic Science Research Program through the National Research Foundation of Korea (NRF) funded by the Ministry of Education (NRF-2021R1I1A1A01050990). B.J. is supported by MnDRIVE (Minnesota’s Discovery, Research, and InnoVation Economy).

S.R. and B.J. designed the study. J.I.H. and J.K. performed the experiments. J.I.H., J.K., S.R., and B.J. analyzed the data. J.I.H., J.K., and B.J. wrote the manuscript. J.I.H., J.K., S.R., and B.J. critically reviewed the manuscript.

We declare no conflicts of interest.

## Notes

### Competing Interest Statement

The authors have declared no competing interest.

