## Supplemental figures for "Phylogenetic Association and Genetic Factors in Cold Stress Tolerance in *Campylobacter jejuni*"

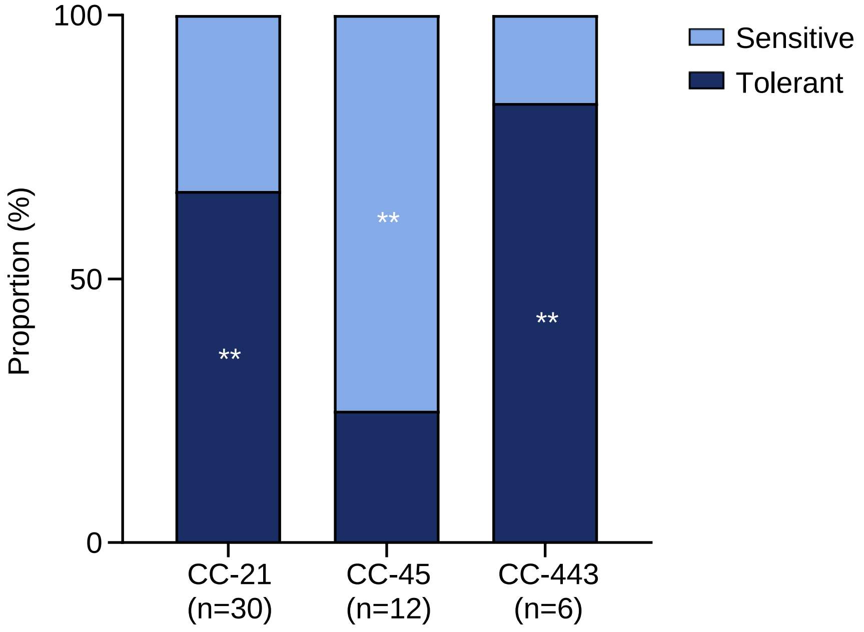


**Figure S1. Distribution of cold stress tolerance of strains among CC-21, CC-45, and CC-443.** The proportions of cold stress-sensitive and cold stress-tolerant strains among CC-21, CC-45, and CC-443 were compared. A chi-square test was conducted for comparison of the proportions of cold stress-tolerant strains in the CCs. $^{**}$, $P < 0.01$; CC, clonal complex.

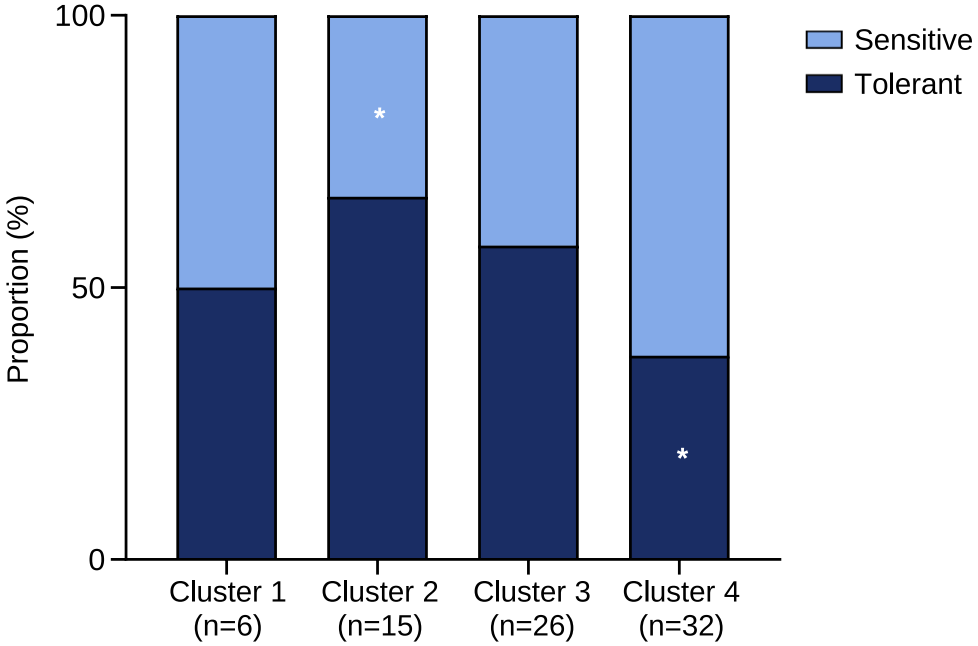


**Figure S2. Distribution of cold stress tolerance of strains belonging to the four Clusters.** The distribution of cold-sensitive strains (n=40) and cold-tolerant strains (n=39) in Cluster 1, Cluster 2, Cluster 3, and Cluster 4 was compared A chi-square test was conducted to compare the proportions of cold stress-tolerant strains in the clusters. *, $P < 0.05$.

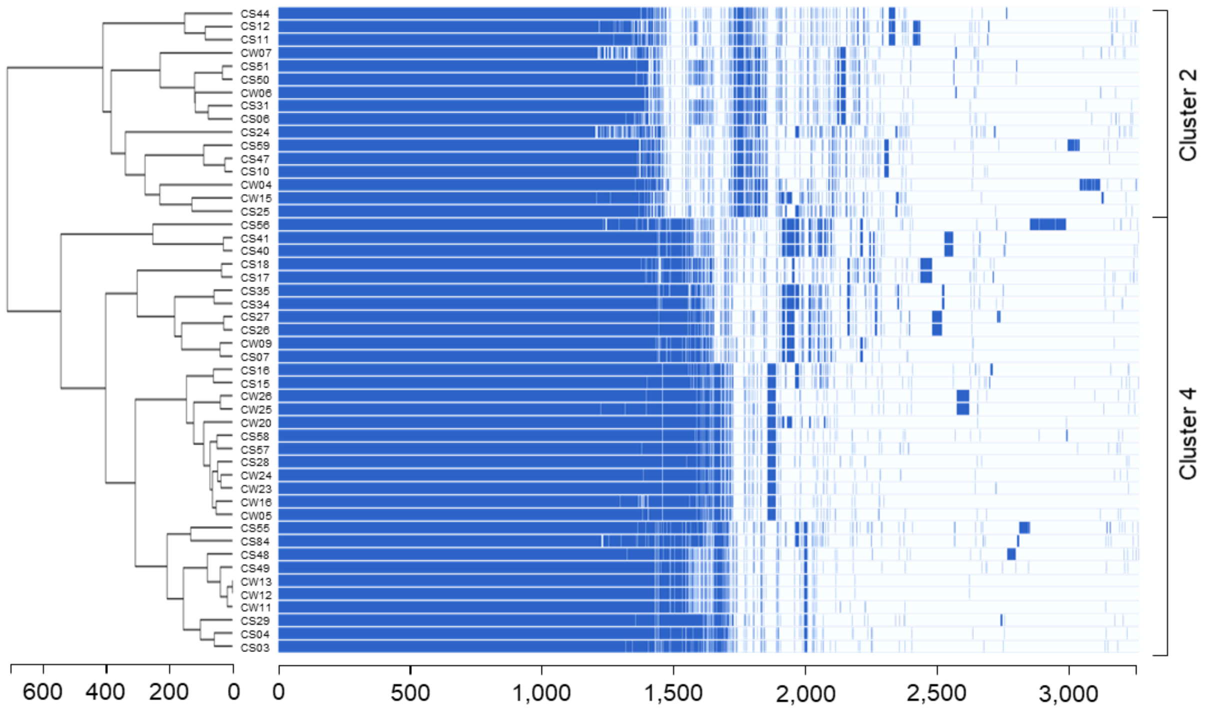


**Figure S3. Pan-genome map comparing cold-sensitive cluster (Cluster 2) and cold-tolerant cluster (Cluster 4).** A pan-genome map was generated with the 47 strains from Cluster 2 (n=15) and Cluster 4 (n=32).

**Table S1. Primers used in this study**

| Primer | Sequence (5′ to 3′) | Reference |
|---|---|---|
| 16s-qPCR-F | ATAAGCACCGGCTAACTCCG | 1 |
| 16s-qPCR-R | TTCCATCTGCCTCTCCCTCA | |
| kan-F | GCGATGAAGTGCGTAAG | 2 |
| kan-R | CGGCTCCGTCGATACTATG | |
| cfrA-Sal1-F | AAAGTCGACGCTAGCAGTTGAAGATTTGCAAA | This study |
| cfrA-BamH1-R | AAAGGATCCCTGCGTTTTCGTTTTCAAATTCTTG | |
| cfrA-inv-F | GGCAAGGTTGAATACAAAGGTGT | |
| cfrA-inv-R | CCAGTAATACCCCTCATAGTGAT | |
| cfrA-comp-Not11-F | TTTGCGGCCGCGGTAAGAGCTGGCTTGTGGA | |
| cfrA-comp-Not1-R | AAAGCGGCCGCGCGTATAGTCGAAGCAGGTAT | |
| pFMBcomC-inv-Not1-F | GCGGCCGCAAGCTAATTGTTTATGGGGAATTAG | |
| pFMBcomC-inv-Not1-R | GCGGCCGCTTATTACTTTGTACTCTAGGGGT | |
| kan-conf-F | CGGGGAAGAACAGTATGTCG | |
| kan-conf-R | CTCCCACCAGCTTATATACCTTA | |
| pFMBcomC-F2 | CCTGTTTCTATGATACCGTGGA | |
| pFMBcomC-R3 | GGGCCTAACAAGACTTGAACTT | |
| cfrA-conf-F1 | AATGCCTACAAAATCAAAGATAGTGATA | |
| cfrA-conf-F2 | AAAGTCCAGGTAAATTCTACAAGAAC | |
| cfrA-conf-F3 | AAAGGATAATGCACCTATTGGTTCTA | |
| cfrA-conf-F4 | TTACAGGCTTTAGAACCCCTTATG | |
| cfrA-conf-R1 | GTGGGGAGGTTCTTGTAGAATT | |
| cfrA-conf-R2 | TTTTCTAGAGAGCCACTCCATGTTTTTCAAG | |
| cfrA-conf-R3 | ATTCCTGGACTTGTGATGGTTCC | |
| cfrA-conf-R4 | CTGCCTTGGCCACTATAACTG | |
| cfrA-P/A-F | TTTGTCGCAGAAGATATTATCTTAGATA | |
| cfrA-P/A-F | TTTGTATCACCCATATAGCGATCTATTT | |
| cfrA-RT-F | CCTGCTACTATCAATGTTATCAC | |
| cfrA-RT-R | CTGACGACGCCCATCAATCA | |

**Table S2. The GenBank accession numbers of the genome sequences of the 79 *C. jejuni* isolates used in the study**

| Sample name | Genome Accession |
|---|---|
| CS01 | JAMGEB000000000 |
| CS02 | JAMGEA000000000 |
| CS03 | JAMGDZ000000000 |
| CS04 | JAMGDY000000000 |
| CS05 | JAMGDX000000000 |
| CS06 | JAMHFV000000000 |
| CS07 | JAMGDW000000000 |
| CS08 | JAMGDV000000000 |
| CS09 | JAMGDU000000000 |
| CS10 | JAMHFU000000000 |
| CS11 | JAMGDT000000000 |
| CS12 | JAMGDS000000000 |
| CS13 | JAMGDR000000000 |
| CS14 | JAMGDQ000000000 |
| CS15 | JAMHFT000000000 |
| CS16 | JAMHFS000000000 |
| CS17 | JAMHFR000000000 |
| CS18 | JAMGDP000000000 |
| CS19 | JAMHFQ000000000 |
| CS22 | JAMGDO000000000 |
| CS23 | JAMGDN000000000 |
| CS24 | JAMGDM000000000 |
| CS25 | JAMGDL000000000 |
| CS26 | JAMGDK000000000 |
| CS27 | JAMGDJ000000000 |
| CS28 | JAMGDI000000000 |
| CS29 | JAMHFP000000000 |
| CS31 | JAMGDH000000000 |
| CS32 | JAMGDG000000000 |
| CS33 | JAMGDF000000000 |
| CS34 | JAMGDE000000000 |
| CS35 | JAMGDD000000000 |
| CS36 | JAMGDC000000000 |
| CS37 | JAMGDB000000000 |
| CS38 | JAMGDA000000000 |
| CS40 | JAMGCZ000000000 |
| CS41 | JAMGCY000000000 |
| CS42 | JAMHFO000000000 |
| CS43 | JAMHFN000000000 |

| | |
|---|---|
| CS44 | JAMHFM000000000 |
| CS45 | JAMHFL000000000 |
| CS47 | JAMGCX000000000 |
| CS48 | JAMGCW000000000 |
| CS49 | JAMGCV000000000 |
| CS50 | JAMGCU000000000 |
| CS51 | JAMHFK000000000 |
| CS52 | JAMHFJ000000000 |
| CS53 | JAMGCT000000000 |
| CS54 | JAMHFI000000000 |
| CS55 | JAMWEZ000000000 |
| CS56 | JAMGCS000000000 |
| CS57 | JAMHFH000000000 |
| CS58 | JAMHFG000000000 |
| CS61 | JAMGCR000000000 |
| CS62 | JAMGCQ000000000 |
| CS63 | JAMHFF000000000 |
| CS64 | JAMHFE000000000 |
| CW01 | JAMGCP000000000 |
| CW02 | JAMGCO000000000 |
| CW03 | JAMGCN000000000 |
| CW04 | JAMGCM000000000 |
| CW06 | JAMGCL000000000 |
| CW07 | JAMGCK000000000 |
| CW08 | JAMGCJ000000000 |
| CW09 | JAMGCI000000000 |
| CW11 | JAMGCH000000000 |
| CW12 | JAMGCG000000000 |
| CW13 | JAMGCF000000000 |
| CW15 | JAMGCE000000000 |
| CW16 | JAMHFD000000000 |
| CW17 | JAMGCD000000000 |
| CW18 | JAMGCC000000000 |
| CW19 | JAMGCB000000000 |
| CW20 | JAMGCA000000000 |
| CW22 | JAMGBZ000000000 |
| CW23 | JAMGBY000000000 |
| CW24 | JAMGBX000000000 |
| CW25 | JAMGBW000000000 |
| CW26 | JAMGBV000000000 |
